# Honokiol inhibits SARS-CoV-2 replication in cell culture

**DOI:** 10.1101/2022.07.26.501656

**Authors:** Clarisse Salgado-Benvindo, Anouk A. Leijs, Melissa Thaler, Ali Tas, Jack L. Arbiser, Eric J. Snijder, Martijn J. van Hemert

## Abstract

SARS-CoV-2 emerged in 2019 and since its global spread has caused the death of over 6 million people. There are currently few antiviral options for treatment of COVID-19. Repurposing of known drugs can be a fast route to obtain molecules that inhibit viral infection and/or modulate pathogenic host responses. Honokiol is a small molecule from *Magnolia* trees, for which several biological effects have been reported,, including anticancer and anti-inflammatory activity. Honokiol has also been shown to inhibit several viruses in cell culture. In this study, we show that honokiol protected Vero E6 cells from SARS-CoV-2-mediated cytopathic effect with an EC50 of 7.8 µM. In viral load reduction assays we observed that honokiol decreased viral RNA copies as well as viral infectious progeny titers. The compound also inhibited SARS-CoV-2 replication in the more relevant A549 cells, expressing ACE2 and TMPRSS2. A time-of-addition assay showed that honokiol inhibited virus replication even when added post infection, suggesting it acts at a post-entry step of the replication cycle. Honokiol was also effective against more recent variants of SARS-CoV-2, including omicron and it inhibited other human coronaviruses as well. Our study suggests that honokiol is an interesting molecule to evaluate in animal studies and clinical trials to investigate its effect on virus replication and pathogenic (inflammatory) host responses.

## INTRODUCTION

Since it emerged in 2019, the Severe Acute Respiratory Syndrome Coronavirus 2 (SARS-CoV-2) lead to a pandemic with a major impact in the whole world. By June 2022 the virus has infected over 500 million people and caused over 6 million deaths globally. SARS-CoV-2 is a member of the betacoronavirus genus within the coronaviridae family, and it is genetically close to SARS-CoV, with almost 80% identity [1]. COVID-19, the disease caused by SARS-CoV-2, often involves mild symptoms like fever, cough, tiredness and loss of taste and/or smell, but it can lead to more serious outcomes, with patients developing shortness of breath, severe pneumonia, respiratory failure or death [2]. Since it emerged in 2019, SARS-CoV-2 had given rise to innumerous variants of interest and/or concern, some of which displayed a worryingly fast spread, increased vaccine escape and/or changes in disease severity [3-5]. Risk factors for severe disease include old age, obesity, and defects in interferon signaling. Host factors are involved in pathogenesis, e.g. via the inflammatory response, but also play a role in viral replication, and therefore constitute interesting therapeutic targets.

Even with the advances in vaccine development, antivirals are still of major importance in order to treat patients that, for a variety of reasons, could not be vaccinated or that did not properly respond to vaccination. Moreover, waning immunity and the continuing emergence of new variants that, to varying degrees, can escape natural or vaccine induced immunity in the population, or the (zoonotic) emergence of yet a new coronavirus makes it even more important to increase our preparedness. By developing antivirals. Preferably not only active against SARS-CoV-2 but to a broad-spectrum of coronaviruses. Compared to vaccines, antivirals also have advantages in terms of storage, distribution, administration and acceptance by part of society. Currently, there are very few options available for treating COVID-19 patients, such as remdesivir, paxlovid and molnupiravir. Issues with costs, route of administration, concerns about side-effects and possible development of resistance regarding the currently approved antivirals against SARS-CoV-2 make it necessary to continue research on potential new antivirals. Repurposing of compounds with already known pharmacokinetic and safety profiles is one way to do this. Early in the pandemic, as part of our drug repurposing efforts, we identified the small molecule Honokiol (HK) as an inhibitor of SARS-CoV-2 replication in cell culture. HK is a polyphenolic lignan compound, extracted from the barks of plants of the Magnolia genus. It has been used in Traditional Chinese Medicine for its analgesic and other effects [6]. In Western medicine HK has also been studied in pharmaceutical and biological studies into its anticancer activities [7-9], anti-inflammatory [10, 11], anti-thrombotic [12], anti-oxidative [13, 14] and antiviral activities [15-17]. HK has been found to modulate several molecular targets, including NF-kB, STAT3, m-TOR and SIRT3 [8, 18-20]. In this study, we focused on the efficacy of honokiol in inhibiting SARS-CoV-2 replication in cell culture. Future studies will assess the effect on the host response to virus, which could play a role in mediating the severity of disease. We found that honokiol decreased replication of the early pandemic more recent variants of SARS-CoV-2, as well as other pathogenic coronaviruses. Honokiol and analogs may be useful in the treatment of SARS-CoV-2 infections and our studies provide a rationale for *in vivo* studies to assess its effect on the infection and pathogenesis linked to the host response to SARS-CoV-2 infection.

## MATERIAL AND METHODS

### 1. Cell culture and compounds and viruses

Vero E6 cells were maintained in Dulbecco’s modified Eagle’s medium (DMEM; Lonza), supplemented with 8% fetal calf serum (FCS; Bodinco), 2 mM L-glutamine, 100 IU/ml of penicillin, and 100 μg/ml of streptomycin (Sigma-Aldrich). A549 expressing ACE2 and TMPRSS2 (here referred to as A549-ACE2-TMPRSS2 cells) were a kind gift from Stuart Neil (King’s College, London, United Kingdom) and are described in [23]. These cells were maintained in DMEM supplemented with 8% FCS, 100 IU/ml of penicillin, 400 ug/mL of G418 (InvivoGen), and 1 ug/mL of Puromycin (Sigma-Aldrich). Huh7 cells were grow in DMEM, supplemented with 8% FCS, 2 mM L-glutamine, non-essential amino acids, 100 IU/ml of penicillin and 100 μg/ml of streptomycin.

The SARS-CoV-2/Leiden-0002 (Genbank: MT510999.1) and SARS-CoV-2/Leiden-0008 (Genbank: MT705206.1) isolates were obtained from nasopharyngeal samples at the LUMC at the first wave of the pandemic. For infections of A549-ACE2-TMPRSS2 cells, a SARS-CoV-2 isolate that was adapted to this cell line was used (Groenewold et al., manuscript in preparation). The SARS-CoV-2 delta variant (Leiden-KUL-Delta1) and Omicron variant (Leiden-O-71084/2021) were isolated at LUMC from a local clinical sample and material kindly provided by the National Institute for Public Health and the Environment (RIVM, Netherlands), respectively. MERS-CoV Jordan-N3 (Genbank: KJ614529.1), SARS-CoV/Frankfurt-1 (Genbank: AY291315.1) and HCoV-229E (Genbank: NC_002645.1) were also used for this study. Infections were done using Eagle’s minimal essential medium (EMEM) with 25mM HEPES (Lonza), supplemented with 2% FCS, 2 mM L-glutamine, 100 IU/ml of penicillin and 100 μg/ml of streptomycin. All experiments with SARS-CoV, SARS-CoV-2 and MERS-CoV were done in the LUMC biosafety level 3 facilities, while HCOV-229E infections were done in a biosafety level 2 laboratory. Honokiol was purchased from MedChem Express as a powder and dissolved in DMSO.

### 2. Cytopathic effect (CPE) reduction assay

Vero E6 cells were seeded in 96-well clusters at a density of 5 × 10^3^ cells/well in 100 uL. Twenty-four hours after seeding, cells were incubated with 2-fold dilutions of compound for 1 hour. After that, half the cells were left uninfected, for analysis of compound’s toxicity, or infected with SARS-CoV-2 at a low MOI of 0.015. After three days, cell viability was measured by MTS assay using the CellTiter 96® AQueous MTS Reagent (Promega). MTS absorbance was measured at 495 nm with an EnVision multiplate reader (PerkinElmer).

### 3. Viral load reduction assay

Cells were seeded at a density of 1 × 10^4^ (Vero E6 and Huh7) or 2 × 10^4^ (A549-ACE2-TMPRSS2) cells per well in 100 uL medium in a 96-well cluster. Twenty-four hours after seeding, cells were treated with increasing concentrations of the compound and were incubated for 6 hours at 37°C. Subsequently, cells were infected for 1 hour with virus at a MOI of 1. Supernatant was harvested at 16 h.p.i. for SARS-CoV, SARS-CoV-2 and MERS-CoV infections, or at 24 h.p.i. for HCoV-229E experiments. To assess viral load, extracellular viral RNA copies were quantified by RT-qPCR, and/or infectious progeny was quantified by plaque assay. The potential cytotoxicity of the compound was always tested in parallel by MTS assay in equally treated, but uninfected, cells.

### 4. Plaque assay

Vero E6 cells were seeded 1 day before infection in regular culture medium at 1.5 × 10^4^ cells/well in 1 mL in 12-well clusters. On the day of the assay, 10-fold serial dilutions of samples were prepared in infection medium. These dilutions were used as inoculum to infect cells for 1 hour at 37°C, after which inoculum was removed and replaced with overlay medium containing 1.2% avicel, 1% antibiotics, 2% FCS and 50mM HEPES in DMEM medium. Cells were incubated at 37°C for 3 days and clusters were fixed with 7.4% formaldehyde. Wells were stained with crystal violet and plaques were manually counted to determine the sample’s infectious virus titer.

### 5. RNA isolation and quantitative real time PCR

RNA was isolated from cell culture supernatants using the Bio-on-Magnetic-Beads (BOMB) method [24] using a Viaflo Assist Plus robotic system (Integra), following sample lysis in a buffer containing 3M guanidine-thiocyanate, 2% N-Lauroyl-Sarcosine sodium salt, 1M Tris-HCl (pH 7.6) and 0.5M EDTA. Equine arteritis virus (EAV) RNA was spiked into the lysis reagent as an internal technical control for RNA isolation efficiency and quality. Viral RNA was amplified by RT-qPCR using the Taqman Fast Virus 1-step master mix (Thermo Fisher Scientific). Primers and probes targeting SARS-CoV-2 RNA-dependent RNA polymerase were described in [25], which were also used for SARS-CoV quantification. MERS-CoV and HCoV-229E primer sets were designed in house, targeting the gene for the nucleoprotein (N) of each virus. For absolute quantification, a standard curve generated from a T7 RNA polymerase *in vitro* transcript containing the necessary RT-qPCR target fragments was used. The reaction was performed in a CFX384 Touch™ Real-Time PCR Detection System (Bio-Rad, Netherlands), with a program of 5 min at 50°C and 20 seconds at 95°C, followed by 45 cycles of 5 seconds at 95°C and 30 seconds at 60°C.

## RESULTS

### 1. Honokiol inhibits SARS-CoV-2 replication in VeroE6 cells

To evaluate if HK could protect cells from SARS-CoV-2 infection, we performed a cytopathic effect (CPE) reduction assay. Infected Vero E6 cells showed increased viability upon treatment with increasing concentrations of HK in a dose-dependent manner, with an EC50 of around 7.8 µM (Fig 1A). In parallel in the same plate, uninfected cells were treated with compound to assess its toxicity. We observed no signs of cytotoxicity in Vero E6 cells for the concentrations tested.

**Figure 1:**
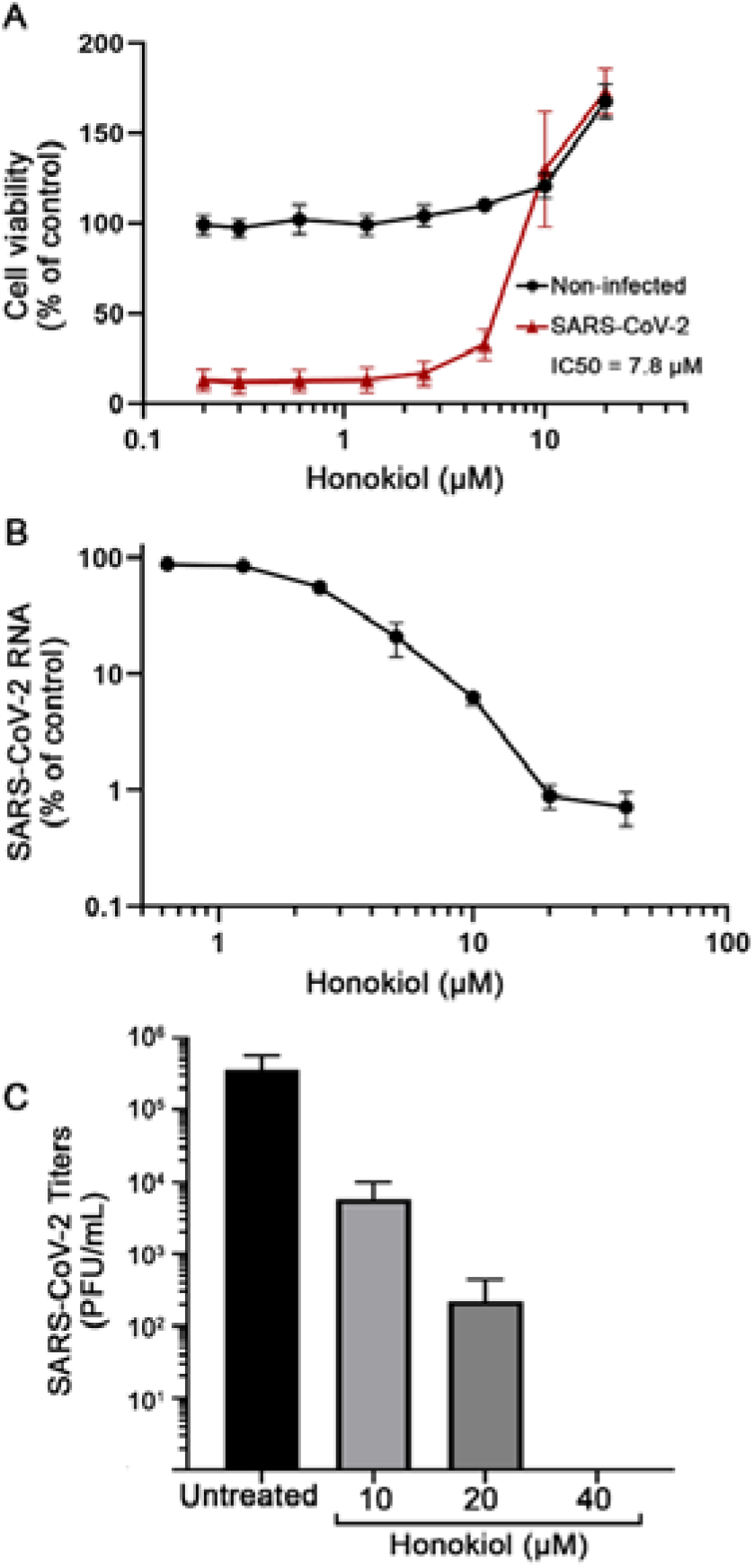
Effect of Honokiol on SARS-CoV-2 mediated cytopathic effect and viral replication in Vero E6 cells. (A) Vero E6 cells were treated with increasing concentrations of HK and then infected with SARS-CoV-2 at an MOI of 0.015. After 3 days, cell viability was measured by MTS assay. The viability of non-infected cells that were treated with compound was determined in parallel to assess cytotoxicity of the compound. (B, C) Vero E6 cells were treated with increasing concentrations of HK and, after 6 hours, were infected with SARS-CoV-2 at an MOI of 1. Supernatant samples were harvested at 16 h.p.i. to quantify SARS-CoV-2 RNA levels by RT-qPCR and infectious progeny titer by plaque assay.

To confirm that the observed protection in the CPE reduction assays was indeed caused by HK inhibition of virus replication, we conducted a viral load reduction (VLR) assay. Vero E6 cells were pretreated with increasing concentrations of HK for 6 hours. After that, cells were infected with SARS-CoV-2 at a MOI of 1 for 1 hour (in the presence of the compound), followed by incubation with medium containing compound. At 16 h.p.i., supernatant was harvested for virus quantification. RT-qPCR showed that HK caused a dose-dependent decrease in viral RNA levels. At 20 µM, a 99% reduction in viral RNA copies, from 1.6 × 10^9^ to 1.3 × 10^7^ copies/mL, was observed (Fig 1B). The infectious virus titer in the supernatant was determined by plaque assays, which showed that treatment with 20 µM of HK caused an approximate 3 log reduction in infectious virus titer (Fig 1C). Cell viability was not affected at these concentrations and only above 40 µM HK starts showing toxicity in VeroE6 cells. This suggested that HK (up to 20 µM) specifically inhibit SARS-CoV-2 replication and protected cells from virus-induced CPE, without causing measurable cytotoxicity.

### 2. Honokiol inhibits SARS-CoV-2 replication at a post-entry step of the replication cycle

To pinpoint which step of the viral replication cycle is inhibited by HK, a time-of-addition assay was performed. Treatment of Vero E6 cells with 20 µM of HK was initiated at different time points, after which it remained present until the end of the assay (unless indicated otherwise). At 0h, cells were infected with SARS-CoV-2 at an MOI of 1 and at 10 h.p.i. supernatant was harvested for quantification of infectious progeny titers. Initially, we assessed the effect of different pre-treatments with 20 µM HK and we did not observe any difference in effectiveness between treatments initiated at any time point between 8 or 1 hour prior to infection (*data not shown*). We then performed assays that involved treatments that were initiated at 1 hour before infection or at 0, 1, 2, 4, 6 or 8 h.p.i. (Fig 2A). Infectious virus titers in the supernatant harvested at 10 h.p.i. were quantified by plaque assays. The maximum effect of the compound was still observed when treatment was initiated as late as 2 hours post-infection. When treatments were started later, HK gradually lost its inhibitory effect, being no longer effective after 8 h.p.i. (Fig 2B). When the compound was only present from 0 to 1 h.p.i. it had no inhibitory effect, suggesting it does not interfere with the early steps of the replication cycle. Together these results suggests that HK acts on a post-entry step of the replication cycle of SARS-CoV-2.

**Figure 2:**
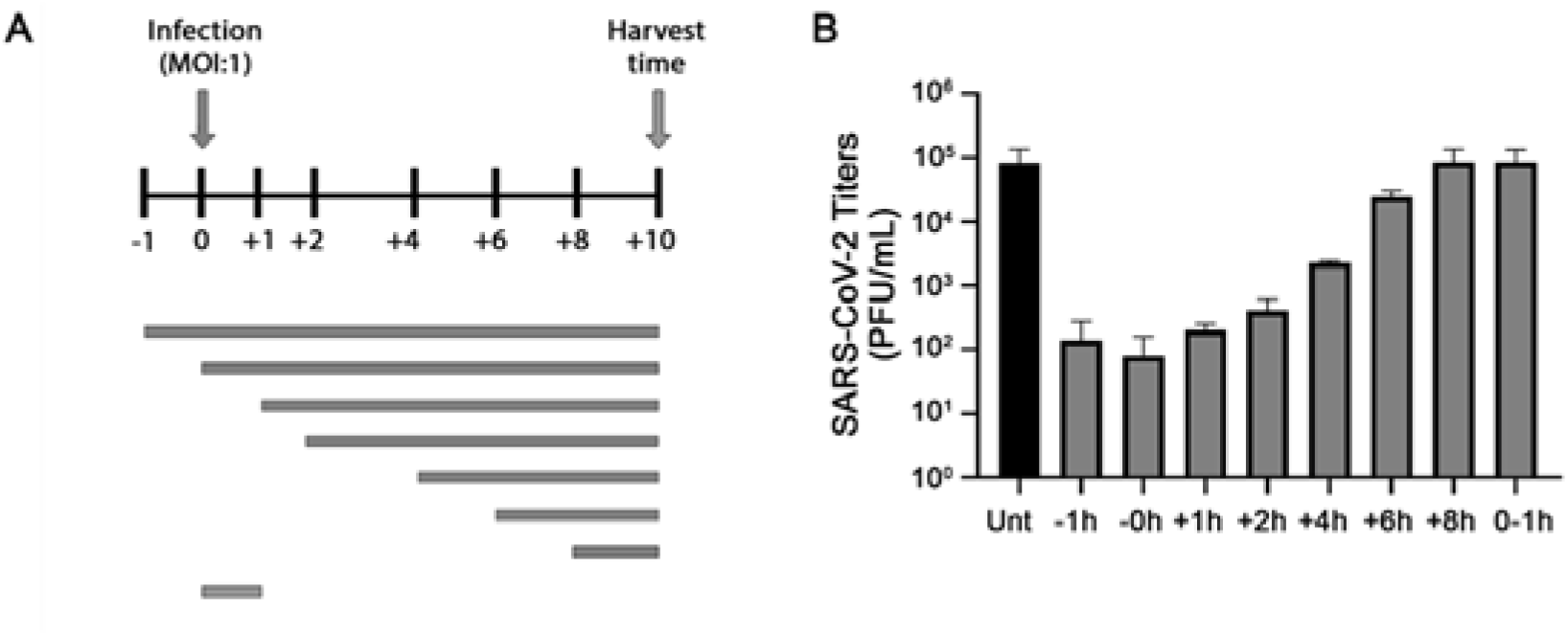
HK inhibited SARS-CoV-2 replication at a post-entry step of the replication cycle. (A) Schematic representation of time-of-addition assay, depicting the different treatment intervals during which infected Vero E6 cells were exposed to 20 µM of HK. (B) At 10 h.p.i., supernatants were harvested and infectious virus titers were determined by plaque assay.

### 3. Honokiol inhibits SARS-CoV-2 replication in human lung cells

To evaluate if HK can also inhibit SARS-CoV-2 in a more relevant cell model, A549-ACE2-TMPRSS2 cells were tested. In a VLR assay, similar to the one described for Vero E6 cells, A549-ACE2-TMPRSS2 cells were pretreated with HK at increasing concentrations and subsequently infected at an MOI of 1. RT-qPCR analysis of supernatant samples harvested at 16 h.p.i. showed that SARS-CoV-2 was also inhibited in this model. Treatment with 10 µM HK already led to a 95% reduction in viral RNA copies, what corresponds to a 2-log decrease in copy numbers (Fig 3A). Cell viability was measured in parallel and showed that, like in Vero E6 cells, 20 µM can be considered a safe nontoxic concentration, whereas at 40 µM toxicity is already detected, with a 50% decrease in cell viability (Fig 3B).

**Figure 3:**
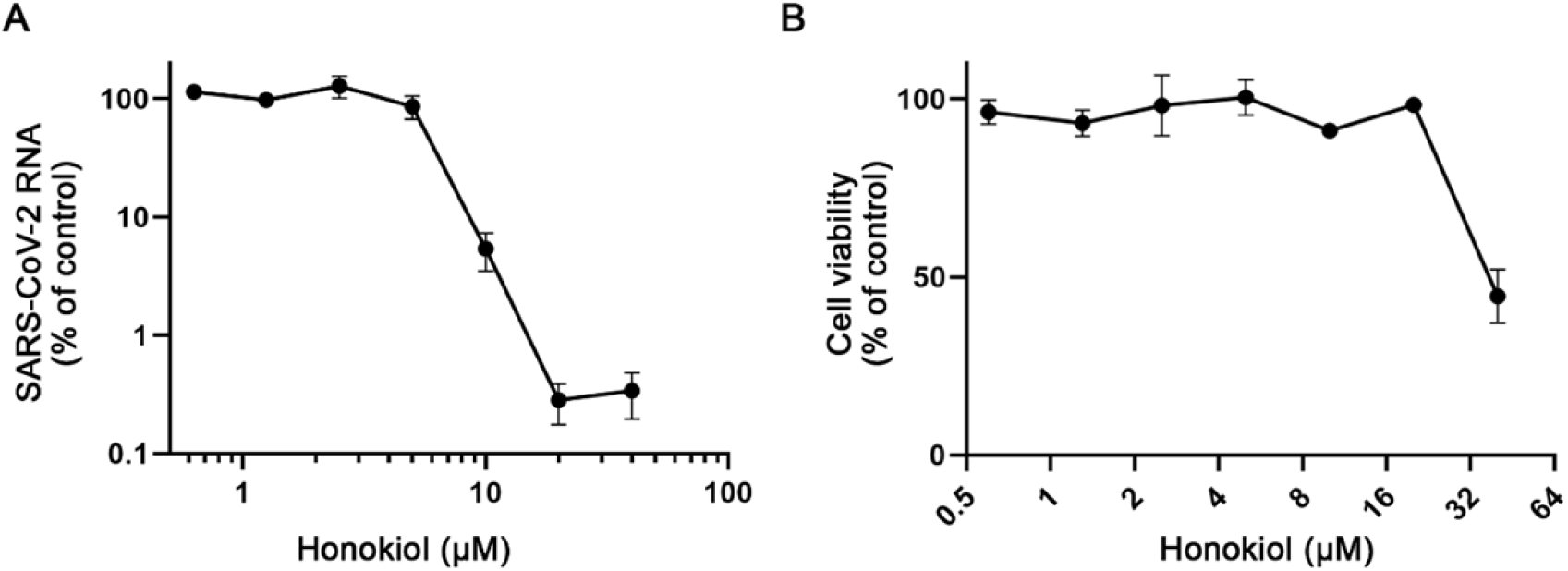
HK inhibited SARS-CoV-2 replication in A549-ACE2-TMPRSS2 cells. (A) A549 cells expressing ACE2 and TMPRSS2 were infected with SARS-CoV-2 at a MOI of 1. Treatment with HK was initiated 6 hours prior to infection and the compound remained present until medium was harvested at 16 h.p.i. to quantify extracellular viral RNA levels by RT-qPCR. (B) compound toxicity was assessed in parallel in uninfected cells treated with HK.

These results show that HK can be a good virus inhibitor even in a more relevant human cell model than the Vero E6 cell, which is a epithelial kidney cell line from African green monkeys. This, besides reenforcing the effectiveness of the compound, also suggests that its antiviral effect is not Vero E6-dependent and can be also observed in other systems.

### 4. Honokiol inhibits replication of a broad spectrum of coronaviruses

To evaluate if HK is effective against other human coronaviruses we studied its effect in Vero E6 cells infected with SARS-CoV or SARS-CoV-2 and in Huh7 cells infected with MERS-CoV or HCoV-229E. Infections were done at an MOI of 1 following a 6-hour pretreatment with HK, after which the compound remained present till the end of the experiment. The medium of MERS-CoV, SARS-CoV, and SARS-CoV-2-infected cells was harvested for virus quantification at 16 h.p.i., and that of HCoV-229E-infected cells at 24 h.p.i. RT-qPCR analysis showed that all viruses were inhibited by HK in a dose-dependent manner (Fig 5A). It is important to note that the maximum nontoxic dose of HK varied depending on the cell line used, with HK showing toxicity in HuH-7 cells already at 20 µM (Fig 5B), while this concentration was not cytotoxic in Vero E6 and A549-ACE2-TMPRSS2. However, also the antiviral effect of HK was observed at lower doses in HuH-7 cells than in the other cells, as Huh-7 cells infected with MERS-CoV or HCOV-229E displayed a more than a 95% reduction in viral RNA copies when cells were treated with only 10 µM of the compound. Together, these data indicates that, despite some differences in its toxicity, HK inhibited coronavirus replication in a cell line independent manner. The compound displayed a broad-spectrum antiviral effect against a range of different pathologically relevant human coronaviruses, i.e. SARS-CoV-2, MERS-CoV, SARS-CoV and HCoV-229E.

### 5. Honokiol is effective against SARS-CoV-2 variants of concern

To investigate if HK was also able to inhibit replication of SARS-CoV-2 variants other than the original early pandemic strain used throughout this study, we tested its efficacy against the two major variants of concern circulating at the time of this project, namely Delta (B.1.617.2) and Omicron (B.1.1.529). Vero E6 cells were treated with HK for 6 hours and were infected with each variant at an MOI of 1. At 16 h.p.i., medium was harvested for determination of the viral load by RT-qPCR targeting the RdRP coding region (Fig 6). At 20 µM, HK inhibited all three variants, showing the strongest effect against the Delta variant, with a ∼2-log reduction in copy number. The early variant was somewhat less sensitive, and the omicron variant was the least sensitive, although still a ∼96% reduction was observed (from 8.6 × 10^7^ copies in untreated cells to 3.3 × 10^5^ copies in cells treated with 20 µM HK).

## DISCUSSION

Honokiol is a small lignan compound that is extracted from the barks, cones and leaves of trees of the Magnolia genus. These plants have been used in Traditional Chinese Medicine to relieve anxiety, depression and pain. In Western medicine anti-inflammatory, anti-thrombotic, anti-oxidative, anti-fungal, anti-arrhythmic and, mainly, anti-tumor properties have been attributed to HK [26]. HK is thought to inhibit tumor progression through modulation of different signaling pathways that behave aberrantly in cancer patients. For example, HK can induce autophagy in different cancer cells by down-regulating the PI3K/Akt/mTOR signaling pathway [8, 27]. HK is also able to down-regulate NF-κB and STAT3 [18, 20], both of which are generally involved in tumor promotion [28]. Other targets of HK include Sirt3 [19], Nrf2 [29], MAPK [30], and SMAD [31] signaling pathways. Some of these factors, such as Nrf2, Sirt3, mTOR [32-34] have been linked to pathways involved in antiviral responses. This provided the rationale for assessing the effect of HK on SARS-CoV-2 infection in cell culture. We demonstrated that in CPE reduction assays HK protected cells from the virus-mediated cytopathic effect in a dose-dependent manner (Fig 1), with an EC50 of approximately 7.8 µM. The effect of HK was cell line-independent as it inhibited SARS-CoV-2 replication in both African green monkey Vero E6 and human A549 cells expressing ACE2 and TMPRSS2 (Fig 2 and 4). Our (single cycle) time-of-addition analysis revealed that HK retains its full inhibitory effect even when treatment is initiated as late as 2 h.p.i., and then gradually loses effectiveness when treatment is initiated later. When the compound was only present during the 1 hour of infection it had no effect. This suggests that HK inhibits a post-entry step of the replication cycle. In contrast to our observations, two previous studies suggested that HK or its analogs can inhibit SARS-CoV-2 infection by targeting the binding and entry steps of the replication cycle. One study suggested that Spike-ACE2 binding was inhibited [35]. However, this study was not performed with infected cells but using artificial assays with pseudotyped viruses or biochemical assays and a less than 50% inhibition was observed at a high dose (50 µM) of HK. Another study [36] suggested that HK inhibits furin-like proteases, but the specificity and efficacy of this inhibition (∼30% at 100 µM) remains debatable considering that a known furin inhibitor was ∼700 times more potent in the same study.

**Figure 4:**
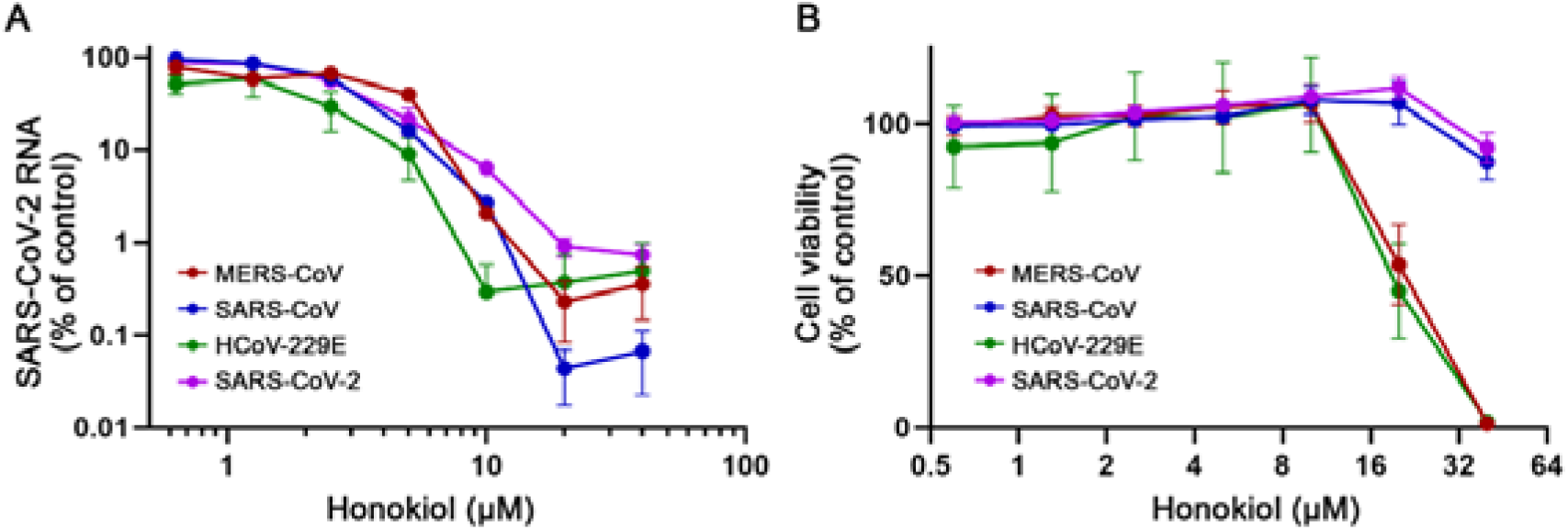
HK inhibited various human coronaviruses. (A) Vero E6 cells were infected with SARS-CoV or SARS-CoV-2 and Huh-7 cells were infected with MERS-CoV or HCoV-229E at an MOI of 1. HK was added 6 hours before infection and remained present till the time of harvest. At 16 h.p.i. (SARS-CoV, SARS-CoV-2 and MERS) or 24 h.p.i. (HCoV-229E), the medium was harvested and the levels of viral RNA were quantified by RT-qPCR. Copy numbers were normalized to the level of untreated infected cells (100%) (B) A viability assay (MTS) was done in parallel to determine the compound’s cytoxicity.

**Figure 5:**
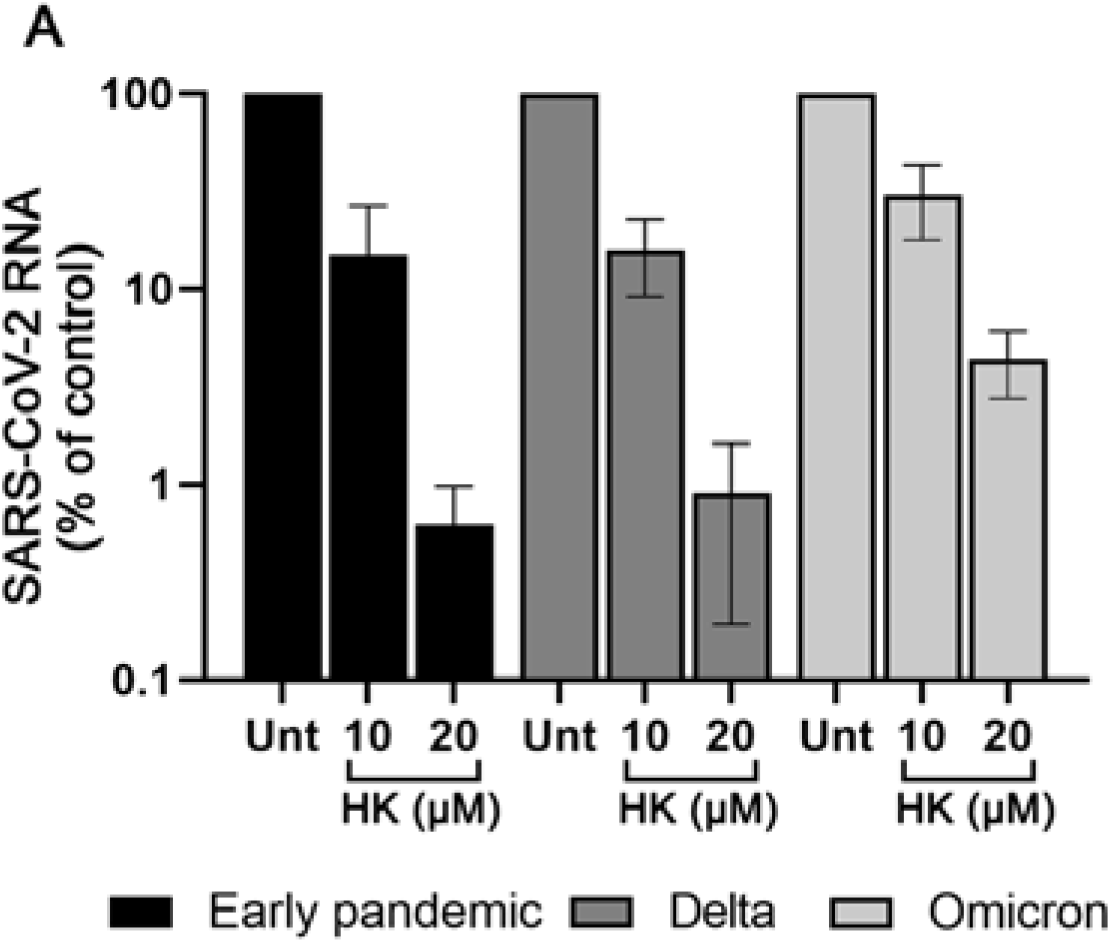
HK inhibited SARS-CoV-2 variants of concern. Vero E6 cells were treated with 10 or 20 µM HK for 6 hours, were infected with a SARS-CoV-2 variants at an MOI of 1, followed by incubation for 16h in the presence of the compound. At 16 h.p.i. supernatant was harvested and RT-qPCR targeting the RdRp gene was used to quantify the extracellular viral RNA levels.

We assessed HK’s spectrum of activity and observed that it also inhibited MERS-CoV, SARS-CoV and HCOV-229E replication in cell culture, in line with the idea that HK likely exerts its antiviral effect through one or more host factors. The fact that HK also inhibited MERS-CoV and HCoV-229E, which use DPP4 and Human aminopeptidase N as receptors, respectively [37, 38], also suggests that it is unlikely that targeting of ACE2 by HK is responsible for the observed antiviral effect. Therefore, we hypothesize that HK inhibits coronavirus replication via one or more host factors involved in the (inflammatory) pathways mentioned earlier, which is currently being studied in more detail.

The massive scale of the SARS-CoV-2 pandemic, complications with (global) vaccine roll out, and the continuing emergence of variants (of concern) that can escape natural or vaccine-induced immunity stresses the importance of developing (multiple) antivirals to increase our preparedness. Direct acting antivirals are a good option, but their spectrum of activity and development of resistance are concerns. Therefore, also compounds that modulate pathways that are involved in the replication of (a broad range of) viruses are interesting candidates to explore as potential antivirals. In particular when this involves repurposing of existing compounds with favorable pharmacokinetics and safety profiles. Besides its antiviral effect against various coronaviruses we have also shown that HK inhibited the two major variants that were circulating at the time of our study, i.e. the delta and omicron variant (Fig 5).

HK appears to be well-tolerated, especially when administered orally, which would increase its acceptance as a therapeutic agent [21]. Studies in mice showed that, following intravenous administration free HK levels in the plasma reached around 200 µg/mL [39]. That would be well above the EC50 that we found in our study (around 5 µg/mL). Moreover, HK has good bioavailability. After a single dose orally administered in healthy rats, HK is rapidly absorbed, reaching its peak plasma concentration in 20 min and reaching various tissues after only 5 minutes. It is slowly eliminated, with a half-life of approximately 290 minutes [21]. After intravenous administration, HK also shows a fast peak followed by elimination, apparently faster than when administered orally (approximately 56 minutes after a 10mg/kg dose) [22]. Some clinical studies have been conducted in humans subjected to HK or whole magnolia bark extract treatments (reviewed in [40]). In one study, three volunteers of the treatment group dropped out due to side effects, while the other 16 subjects completed the study without any signs of serious adverse events [41]. Other studies did not report adverse effects related to the treatments.

Effective treatment of SARS-CoV-2 infection will require both antiviral agents as well as agents that modify the host (inflammatory) response. Human aging is associated with a more severe inflammatory response to SARS-CoV-2, and Sirt3 activators such as honokiol have anti-inflammatory effects in vivo, which could have the additional benefit of reducing pathologic inflammation. In conclusion, the safety profile of HK, plasma levels that can be reached after administration and its broad-spectrum antiviral effect against multiple coronaviruses and possible effect on inflammation make it an interesting compound to explore in animal studies and clinical trials as (part of) treatment for SARS-CoV-2 infections.

## ACKNOWLEDGMENTS

The authors would like to thank Jessika Zevenhoven, Patrick Wanningen, Mirjam Groenewold and Mariska Huizen for technical assistance and Stuart Neil (Kings college, London) for providing the A549-ACE2-TMPRSS2 cell line. Clarisse Salgado-Benvindo was supported by the Coordination for the Improvement of Higher Education Personnel (CAPES) (Process nr. 88881.171440/2018-01), Ministry of Education, Brazil.

